# Weather and leaf age separately contribute to temporal shifts in phyllosphere community structure and composition

**DOI:** 10.1101/2024.06.21.600104

**Authors:** Jacob A. Heil, Allison Simler-Williamson, Miranda L. Striluk, Danielle Trawick, Rachel Capezza, Chadwick DeFehr, Aubrey Osorio, Bruce Finney, Kathryn G. Turner, Leonora S. Bittleston

**Affiliations:** ¹Department of Biological Sciences, Boise State University, Boise, Idaho, USA; ²Department of Biological Sciences, College of Western Idaho, Nampa, Idaho, USA; ³Department of Biological Sciences, Idaho State University, Pocatello, Idaho, USA

**Keywords:** microbial community, phyllosphere, Artemisia, leaf microbiome, fungi, community ecology, sagebrush steppe, sagebrush, time series

## Abstract

Microbial communities living on plant leaves can positively or negatively influence plant health and, by extension, can impact whole ecosystems. Most research into the leaf microbiome consists of snapshots, and little is known about how microbial communities change over time. Weather and host physiological characteristics change over time and are often collinear with other time-varying factors, such as substrate availability, making it difficult to separate the factors driving microbial community change. We leveraged repeated measures over the course of an entire year to isolate the relative importance of environmental, host physiological, and substrate age-related factors on the assembly, structure, and composition of leaf-associated fungal communities. We applied both culturing and sequencing approaches to investigate these communities, focusing on a foundational, widely-distributed plant of conservation concern: basin big sagebrush (*Artemisia tridentata* subsp. *tridentata*). We found that changes in alpha diversity were independently affected by the age of a community and the air temperature. Surprisingly, total fungal abundance and species richness were not positively correlated and responded differently, sometimes oppositely, to weather. With regard to beta diversity, communities were more similar to each other across similar leaf ages, air temperatures, leaf types, and δ^13^C stable isotope ratios. Nine different genera were differentially abundant with air temperature, δ^13^C, leaf type, and leaf age, and a set of 20 genera were continuously present across the year. Our findings highlight the necessity for longer-term, repeated sampling to parse drivers of temporal change in leaf microbial communities.

**Open Research Statement:** All ITS DNA amplicon sequence raw data are deposited in the NCBI Sequence Read Archive (SRA), BioProject number PRJNA1107252, data will be released upon publication. All community data, metadata, taxonomic data, and R code necessary to reproduce these results are deposited in the GitHub repository archived on Zenodo: 10.5281/zenodo.11106439.

## Introduction

The phyllosphere community, here defined as the group of organisms living internally and externally on leaves, influences plant health through positive and negative interactions, such as pathogenicity, growth promotion, nutrient acquisition, disease mediation, and more (Leveau 2019; Stone, Weingarten, and Jackson 2018). All plants harbor phyllosphere communities and the impact of these communities on host plant health have implications for the functioning of whole ecosystems (e.g., plant community structure, soil quality, and biogeochemical cycling) (Leveau 2019; Laforest Lapointe and Whitaker 2019). Because of the potentially outsized influence phyllosphere communities have on ecosystems, it is essential to understand the factors influencing their assembly, composition, and structure.

Multiple stochastic and deterministic processes shape microbial community structure and composition over time (Nemergut et al. 2013). Stochastic processes include priority effects and drift caused by birth, death, colonization and extinction (Nemergut et al. 2013; Zhou and Ning 2017). On the deterministic side, spatial factors (e.g., distance, landscape heterogeneity) affect the ability of phyllosphere community constituents to disperse and colonize (Fukami 2015). Weather and other abiotic variation may drive both selective processes (e.g., growth or mortality due to temperature or moisture), as well as influencing the ability of organisms to disperse (e.g., via wind direction and speed). Finally, the host plant itself influences its microbiome through selective factors such as plant chemistry, as well as affecting dispersal of organisms through the plant’s physical structure (Stone and Jackson 2019; Xueliang et al. 2020). Despite our knowledge of these ecological drivers, little is known about how assembly processes of phyllosphere community composition shift in relative importance over time (Dini-Andreote et al. 2015).

Isolating the relative importance of factors driving community assembly over time is difficult in observational designs. Weather measurements are strongly collinear with long-term climate and time-invariant site factors (e.g., soils, population structure). Thus, linking short-term weather events to ecological responses requires either simplifying assumptions (e.g., averaging of weather variables over large windows) or repeated temporal sampling (Teller et al. 2016), which is often omitted from many study designs (e.g., in plant ecology; Vaughn & Young, 2010 (Vaughn and Young 2010), Werner et al., 2020 (Werner et al., n.d.)). Few studies have examined how temporal change affects the structure and diversity of phyllosphere communities (e.g., (Barge et al. 2019; Almario et al. 2022; Dove et al. 2021; VanWallendael et al. 2022; Wagner, Busby, and Balint Kurti 2020)). Barge et al. (Barge et al. 2019) describes variation in samples of the *Populus trichocarpa* phyllosphere taken a year apart. Copeland et al. (Copeland et al. 2015) qualitatively observed shifts in community structure in response to heavy precipitation events and found correlations between the relative abundance of some highly abundant taxa and sampling date, developmental stage, and rainfall. It also remains unknown how quickly phyllosphere community structure and diversity change, although there is some evidence for large shifts in diversity in less than a week (Copeland et al. 2015; Maignien et al. 2014). To address these knowledge gaps, we sampled phyllosphere communities every two weeks over one full year from on leaf substrates with differing ages, to experimentally isolate the relative influence of community age, weather conditions, and host physiology on phyllosphere community change.

Leaf-associated communities are different in structure and diversity from the communities living in soil, stems, air, etc. (Müller et al. 2016). New leaves represent uncolonized substrates for microbial dispersal and community formation (Schlecter, Miebach, and Remus-Emsermann 2019). Some past phyllosphere studies have observed community change over the course of the entire lifespan or the entire growing season of a plant in annuals or perennials with temporary foliage (Almario et al. 2022; VanWallendael et al. 2022; Copeland et al. 2015). For deciduous plants, microbes must disperse onto new leaves annually from sources including air, soil, stems, or from inside the plant (Vacher et al. 2016), however, evergreen plants allow for vertical transfer of more similar communities to new leaves and continuously maintain leaf specialist species throughout time.

Our study leverages the unique features of leaf development in basin big sagebrush (*Artemisia tridentata* subsp. *tridentata*, hereafter, sagebrush) to understand drivers of temporal phyllosphere community change. Sagebrush is the foundation species in the sagebrush steppe ecosystem, which covers the majority of the Western United States (Crist et al. 2019). In this ecosystem, sagebrush species influence the structure of plant and animal communities by serving as nurse plants, physical habitat, and an essential food resource (Davies et al. 2011; Olsoy et al. 2020). A few investigations of sagebrush’s rhizosphere microbes exist (e.g., Gehring et al., 2016 (Gehring et al. 2016)); however, the sagebrush phyllosphere remains undescribed. Beyond its ecological importance, this species is an excellent focal plant for researching leaf microbial communities over time because it maintains functional foliage throughout the entire year (Miller and Shultz 1987) in the form of persistent (late Summer-Spring) and ephemeral (Spring-Summer) leaves, providing a continuous habitat for leaf microbe specialists and allowing the partitioning of weather effects from the estimated age of leaves.

Here we report results from whole-community internal transcribed spacer (ITS) amplicon sequencing of sagebrush leaves every two weeks for an entire year, along with quantitative PCR to determine absolute abundances, and microbial culturing to separate epiphytes from endophytes and to map DNA sequencing data to living organisms. Using leaf isotope data collected at each time point and a full year of hourly weather data, we modeled fungal community diversity as a response to these factors to isolate the relative effects of estimated leaf age, weather, and host plant physiology on phyllosphere communities. We also identified genera that were differentially abundant and those that were continually present at detectable abundances. Results suggest that diversity changes over time and in short intervals, that changes in diversity correlate with age of a community, air temperature, precipitation, and leaf type, that a small number of species consistently persist in the phyllosphere, and that different species are dominant under different weather conditions.

## Materials and Methods

### Phyllosphere community collection

We collected leaf samples from a stand of sagebrush adjacent to the Lower Weather Station (43.6885278, -116.16991) operated by Boise State University Department of Geosciences in the Dry Creek watershed of the Boise foothills in southwestern Idaho (McNamara 2018). Beginning on March 12, 2021, we collected samples every two weeks for the duration of a year (26 time points), ending on February 24, 2022. We collected our samples from the same 6 plants at each time point (Figure 1).

**Figure 1.**
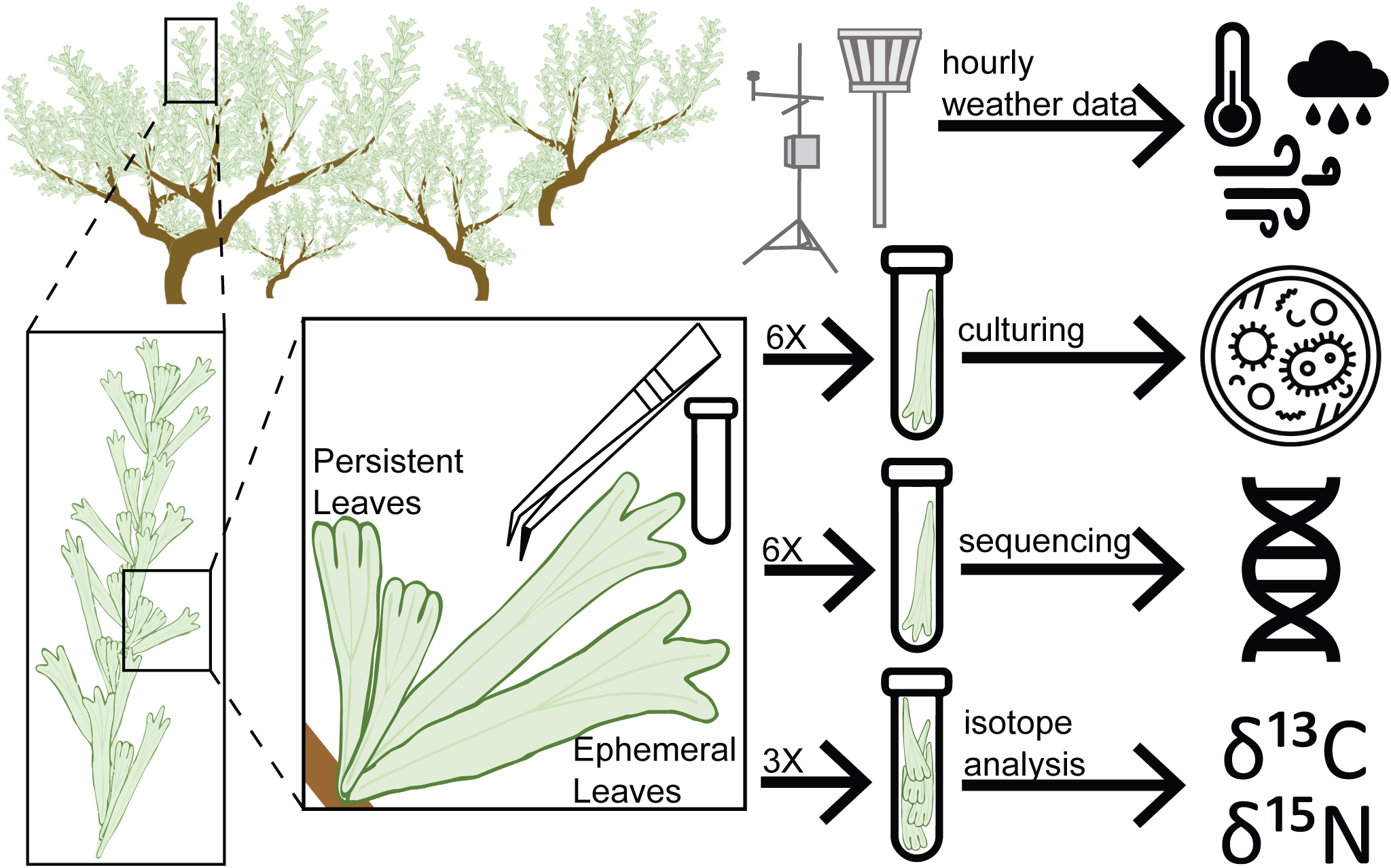
Overview of methods. We collected samples from a single stand of sagebrush in parallel with hourly weather data from a weather station at the site. From each of six plants we collected six individual leaves for culturing of endophytes and epiphytes, six individual leaves for sequencing, and three tubes filled with leaves for isotope analysis.

To collect the leaves, we used sterile field technique: we wore nitrile gloves and collected leaves with metal forceps, sterilizing gloves and equipment with 95% ethanol in between each leaf collected. We collected leaves in sterile microcentrifuge tubes, placing one leaf in each tube intended for microbiome analysis and collecting three tubes filled with leaves for stable isotope analysis. Leaves were collected at different parts of the sagebrush canopy to generally represent the whole-plant leaf community. We collected leaves for two parallel data sets: culturing and sequencing. For each approach we collected six leaves from each of six plants, totaling 36 per sample date, with the exception of four dates when both leaf types were present and we collected 36 of each type, totaling 72 (see supplement for more details). In total we collected 792 persistent leaves and 288 ephemeral leaves each for both culturing and sequencing. Samples for sequencing were stored on ice during transport and placed in -80°C until downstream analysis.

### DNA Sequencing and qPCR

See the supplement for information about the isolation and barcoding of fungal morphospecies. For amplicon sequencing, DNA was extracted from a composite sample of three leaves per plant at each sampling date using the ZymoBIOMICS 96 MagBead DNA Kit (Zymo Research Corp, Irvine, CA, USA; Cat # D3408) and evaluated DNA quality using the AccuClear Ultra High Sensitivity dsDNA Kit (Cat # 31028). Amplicon library preparation and sequencing were done at the Molecular Research Core Facility at Idaho State University (RRID:SCR_012598). We amplified the ITS1 region of the fungal rDNA gene using the ITSf1/ITS2 primer pair (Gardes and Burns 1993) . We checked the quality of our sequences, removed chimeras, and generated amplicon sequence variants (ASVs) using QIIME2 version 2022.2 (Bolyen et al. 2019) and the DADA2 plugin (Callahan, McMurdie, and Rosen 2016), resulting in a data set of 4,998 ASVs and 179 samples, after removing one outlier with drastically higher abundance counts than our other samples. See the supplemental information for details about taxonomic assignment. To determine absolute abundance of the individual ASVs in each sample we calculated the concentration of each sample with quantitative PCR (qPCR) using the Zymo Research Femto Fungal DNA Quantification Kit (Cat # E2007), determined the total copy number of ITS amplicons in sample, and multiplied the copy number per sample by the proportion of each ASV in sample.

### Weather and leaf-level covariates

To determine the effect of different factors on the structure of the sagebrush phyllosphere community, we collected data on estimated leaf age (the time since a leaf type has emerged, hereafter ‘leaf age’), weather, and host plant physiology. We calculated the estimated leaf age by marking the time that we first observed and collected a leaf type as time zero and added one for each preceding time point where we collected that leaf type. This leaf age measurement allowed us to test for changes in a community from emergence of new leaves, when they are new substrates for microbial communities to colonize, to leaf senescence. We also collected hourly weather data from the adjacent BSU Dry Creek Lower Weather Station for the duration of the experiment including air temperature, wind speed, precipitation, UV intensity, snowfall, and various measurements of soil moisture and temperature (Appendix S1: Figure S1). We also recorded host plant identity and leaf type (ephemeral or persistent), using the visual guidelines listed in the supplemental information.

We measured leaf carbon (δ^13^C) and nitrogen (δ^15^N) stable isotopes for each plant as a metric of seasonal variation in host photosynthesis and metabolism associated with drought stress. We dried *A. tridentata* leaves at 60°C for 48 hours and ground them with a Wig-L-Bug (International Crystal Laboratories, Garfield, NJ, USA). We encapsulated and analyzed approximately 3.5 mg of dry sample for stable isotope ratios at the Idaho State University Stable Isotope Laboratory using a Costech ECS 4010 elemental analyzer interfaced with a Thermo Delta V Advantage continuous flow isotope ratio mass spectrometer (Thermo Fisher Scientific, Waltham, MA, USA). All stable isotopic data are reported in standard delta notation (δ¹³C, δ¹ N) relative to the Vienna PeeDee Belemnite and atmospheric N_2_ reference standards. Analytical precision, calculated from analysis of standards distributed throughout each run, was ≤±0.2‰ for both C and N stable isotopes, and <±0.5% of the sample value for %N and %C.

### Analysis of seasonal shifts in diversity and richness

We analyzed our data in the R program version 4.2.3 (Team 2023). We developed Bayesian generalized linear mixed models (glmms) of phyllosphere diversity, based on the causal assumptions of a directed acyclic graph (DAG), designed to isolate the effects of leaf age (separate from type), weather, and host physiology (Figure 2b). Our DAG assumes that leaf age, weather, and host physiology all covary with time, and that plant identity may additionally influence diversity, microclimatic variation, and time-invariant aspects of host physiology. Given that plants were described by site-level weather and sampled similarly for ephemeral and persistent leaves and were located within a single block, we do not assume links between plant identity and leaf age or weather.

**Figure 2.**
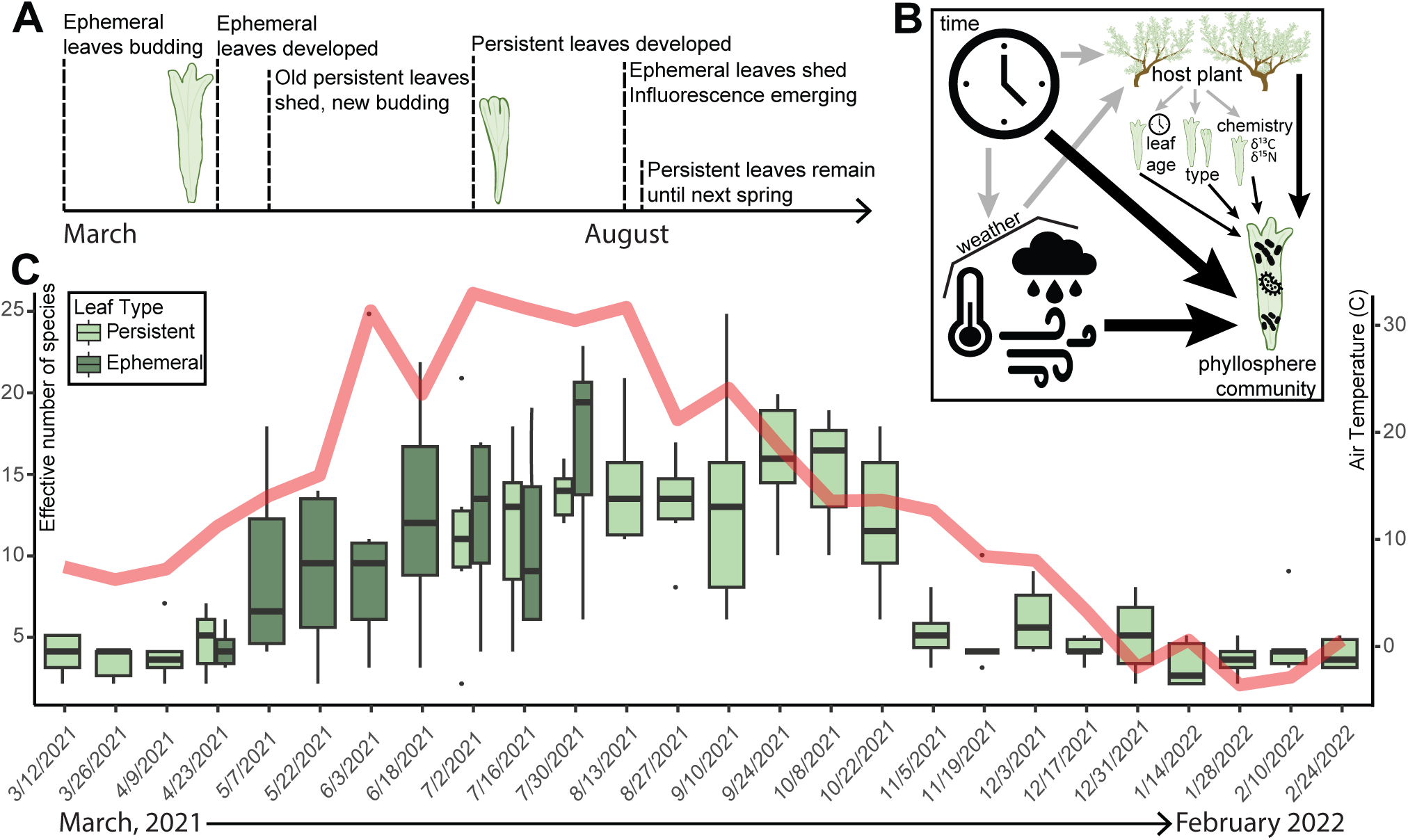
Fungal diversity changes over time and with leaf type. (a) Sagebrush exhibits an annual succession of leaf types, allowing for the isolation of effects of weather variation and leaf age over time. Ephemeral leaves emerge in early spring, mature by late spring, and senesce by late summer. Persistent leaves emerge in late spring, mature in summer, and senesce the next spring. (b) Directed acyclic graph (DAG) of our hypothesis of the sagebrush phyllosphere system. Black arrows are a direct effect and grey arrows are indirect. We hypothesize that time influences phyllosphere communities directly and indirectly via its effects on the host plant and weather. We hypothesize that weather has a direct effect on the phyllosphere community and an indirect effect through its effect on the host plant physiology. The host plant directly influences phyllosphere communities via the age of leaves as community substrates, leaf chemistry, and physiological differences between leaf types. (c) We observed a significant change in ENS over time (p < 0.001) sometimes within our two-week sampling window. Boxplots are colored by leaf type (see legend) and the red line indicates average temperature (°C) at our site.

Based on this DAG, we fit negative binomial glmms for response variables (ASV richness, abundance, and effective number of species) using the brms package and weakly informative priors (Bürkner 2017). Each model contained: 1) leaf age (in days, measured as time since first collection) to describe shifts in community composition associated with substrate age and colonization; 2) variables describing mean air temperature, mean wind speed, and total precipitation for the two week interval preceding each sampling point, to describe the effects of various weather drivers; and 3) leaf δ¹³C and δ¹ N, to describe seasonal variation in host physiology, associated with photosynthetic activity and drought stress. We included a quadratic term for mean air temperature and leaf age, to model possible upper thermal limits for fungi and hump-shaped relationships between time and diversity, respectively. Models also included a varying intercept for plant identity, to account non-independence of repeated measures in its error structure. All continuous variables were centered and scaled by two standard deviations.

All glmms were constructed with the brms package (Bürkner 2017), which uses a Markov Chain Monte Carlo (MCMC) sampler. Models employed brms’ standard (weakly informative) priors, burn-in period, and chain specifications. We checked effective sample size and model convergence, indicated by Gelman–Rubin statistics close to 1 and stable, well-mixed chains (Gelman and Rubin 1992). For additional details on our alpha diversity models, please see the supplement.

### Analysis of microbial community composition and differential abundances

We calculated absolute abundance per ASV, determined total sample abundance, and determined ASV richness. We used the *vegan* package (Oksanen et al. 2022) to calculate diversity metrics. We measured Effective Number of Species (ENS) by taking the exponent of Shannon diversity using the *diversity* function, calculated community dissimilarity using Bray-Curtis with the *vegdist* function, ordinated the dissimilarity data using NMDS with the *metaMDS* function, and calculated differences in beta diversity by first performing a distance based redundancy analysis (dbRDA) with the *dbrda* function with Bray-Curtis dissimilarities as our response variable and the same model formula for the explanatory variables as in our alpha diversity model, controlling for plant identity by using “condition.” We used *anova.cca()* to assess statistical significance of these included constraints in our dbRDA.

We fit differential abundance analyses using the package ANCOMBC (Lin and Pedadda 2020) using the same variables as our alpha and beta diversity models, controlling for plant identity by using “group.” We used the *ggplot2* (Wickham 2016), *effects* (Fox and Weisberg 2018), and *ggcorrplot* (Kassambara 2022) packages to visualize results.

## Results

### Culture-based patterns in phyllosphere composition and structure

From our culture collection, we barcoded a subset of all cultured morphospecies and identified 29 fungal genera and one bacterial genus (*Bacillus*) with Sanger sequencing. We did not comprehensively sequence every morphospecies at every sample date, but noted if they were epiphytes or endophytes as well as their richness, abundance, and physical characteristics. Morphospecies richness was significantly positively correlated with ASV richness from our sequences (r = 0.219, p = 0.003). Abundance was not significantly correlated between sequencing and culturing (p = 0.104). From our culturing results, we observed qualitatively that three of the most consistently present and abundant morphospecies were from the genera *Aureobasidium*, *Filobasidium*, and *Cladosporium*. Endophytes recovered from culturing represented less abundant genera that were consistently endophytic, including: *Coniochaeta, Penicillium,* and *Preussia*. Of these, only *Preussia* was found to be continually present in the phyllosphere community.

### Sequencing-based patterns in phyllosphere alpha diversity

We found that all alpha diversity measures (ASV richness, ASV abundance, and effective number of species) varied significantly over time, sometimes with large changes occurring during the two weeks between samples (Figure 2c, Appendix S1: Figure S2, Appendix S1: Table S1). ASV richness and ENS were positively correlated with each other (r = 0.625, p < 0.001), which we would expect since they are very similar measures, with ENS just adding an element of evenness. Surprisingly, both were negatively correlated with total abundance (ASV richness: r = -0.334, p = 5.064e-06; ENS: r = -0.224, p = 0.003).

### Effects of leaf age and weather on ASV richness

After accounting for differences in leaf type, leaf age exhibited non-zero linear and quadratic relationships with ENS and ASV richness (Appendix S1: Table S2, Figure 3e), separately from leaf type, weather, and leaf physiology. Holding the value of other variables at their means, the ENS increased with leaf age to a maximum of 10.96 species at 16 weeks and then decreased to a minimum of 4.61 species at 42 weeks (Figure 3a). Mean ASV richness increased with leaf age to a maximum of 72 ASVs at 20 weeks and then decreased to a minimum of 38 ASVs at 42 weeks (Figure 3a).

**Figure 3.**
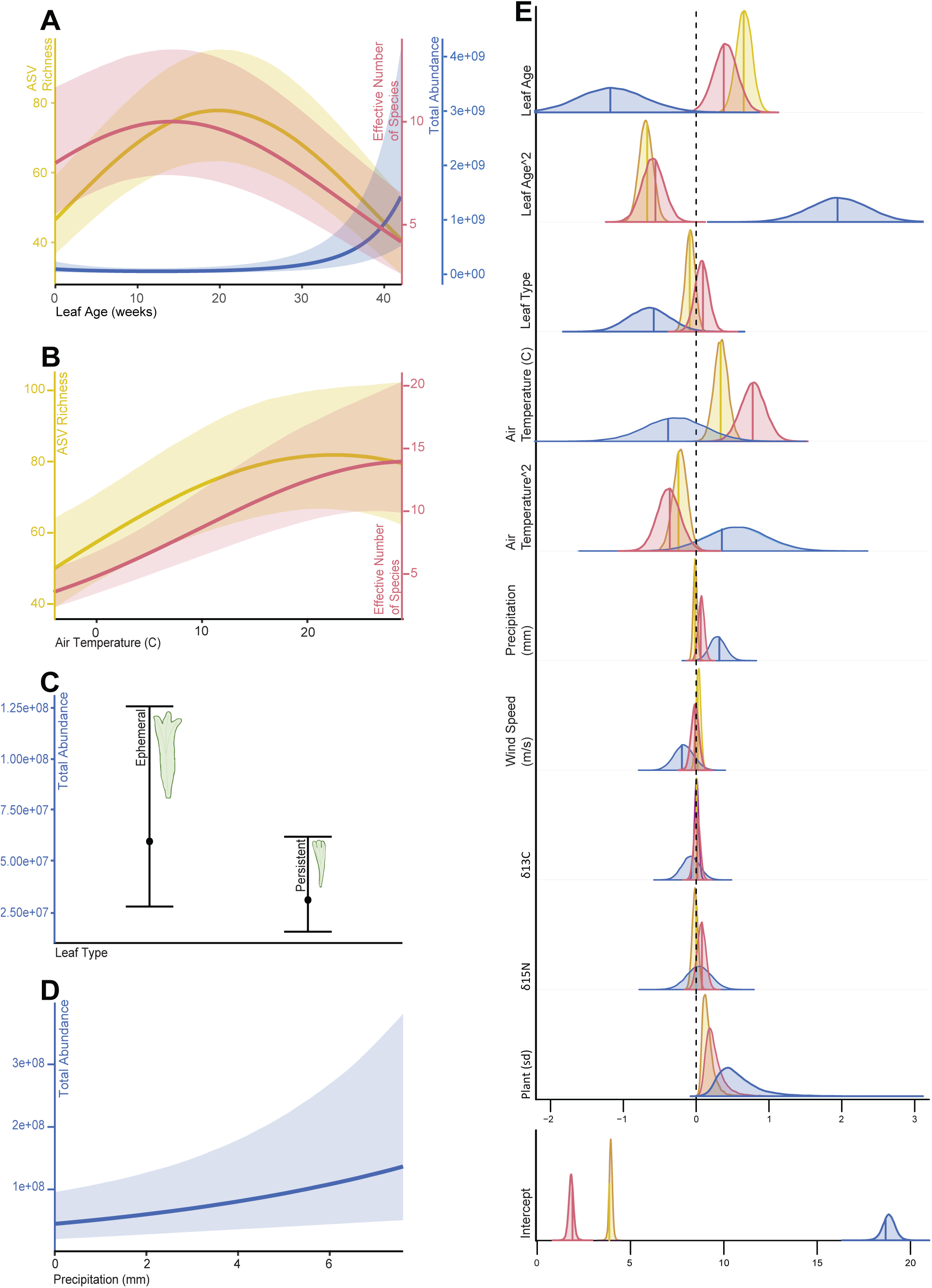
Our model highlights how fungal richness and abundance change with host and environmental factors. (a-d) Marginal effects of leaf age (a), air temperature (b), leaf type (c), and precipitation (d) on ASV richness (yellow), ENS (red), and total abundance (blue), while holding other variables at their means. Plots only depict relationships for variables with 95% credible intervals that do not contain zero. (e) Posterior parameter distributions for scaled variables on ASV richness (shape estimate = 13.61), total abundance (shape estimate = 0.69), and ENS (shape estimate = 11.37), with median estimates shown in solid lines and shaded 95% credible interval regions. Distributions in the “plant” panel are standard deviations from varying intercept for plant identity.

After accounting for the effects of leaf age, type, and physiology, linear and quadratic terms for air temperature had nonzero relationships with ASV richness and ENS (Appendix S1: Table S2, Figure 3e). Mean ASV richness increased with air temperature from a minimum of 46.33 ASVs at -3.99° C to a maximum of 80.96 ASVs at 20.42° C, before declining at greater temperatures (Figure 3b). Mean ENS was predicted to increase with air temperature from a minimum of 3.83 species at -3.99° C to a maximum of 15.32 species at 27.72° C, while holding all other variables at their means (Figure 3b). The 95% credible intervals for the effects of all other included variables on ASV richness and ENS each contained zero.

### Effects of leaf age and weather on microbial abundances

We found that total abundance was most strongly related to leaf age, leaf type, and precipitation. We observed an inverse response to leaf age and air temperature compared to ASV richness and ENS. Leaf age had a negative non-zero relationship with total abundance (Appendix S1: Table S2; Figure 3e). Holding all other variables at their mean value, our model predicted that abundance initially decreased with leaf age to a minimum of 3.29 × 10^7^ at 6 weeks and then increased to a maximum of 1.43 × 10^9^ at 42 weeks (Figure 3a). Ephemeral leaves were more likely to have higher abundance (Appendix S1: Table S2; Figure 3c,e). The effect of air temperature on abundance was opposite to the effect we observed on ASV richness and ENS. Air temperature had a generally negative and quadratic relationship with abundance (probability of direction = 76%), though the 95% credible intervals contained 0. Total abundance was lowest at 4.86°C; it increased from 2.6520893e7 to 1.00731505e8 across the range of precipitation values observed in our study (Appendix S1: Table S2; Figure 3d). The 95% credible intervals for the effects of all other included variables on abundance contained zero.

### Beta diversity

### Effects of leaf age, weather, and host plant factors on Beta diversity

We found that community similarity across samples was influenced by a combination of leaf age, weather, and host plant factors (Appendix S1: Table S2). Based on dbRDA, we found that dissimilarity in community structure was significantly correlated with leaf age (F = 1.638, p = 0.007, Figure 4) and its quadratic term (F = 3.145, p = 0.001). Likewise, dissimilarity in community structure was significantly correlated with air temperature (F = 2.743, p = 0.001) and its quadratic term (F = 2.162, p = 0.003), as was leaf δ^13^C values (F = 2.162, p = 0.001, Figure 4). Lastly, leaf types had significantly different community composition from each other (F = 1.608, p = 0.013, Figure 4), while precipitation (F = 0.982, p = 0.452), wind speed (F = 1.040, p = 0.334), and δ^15^N (F = 1.146, p = 0.192) were all nonsignificant. Together, these variables explained only 12.3% of the variation across all of our samples.

**Figure 4.**
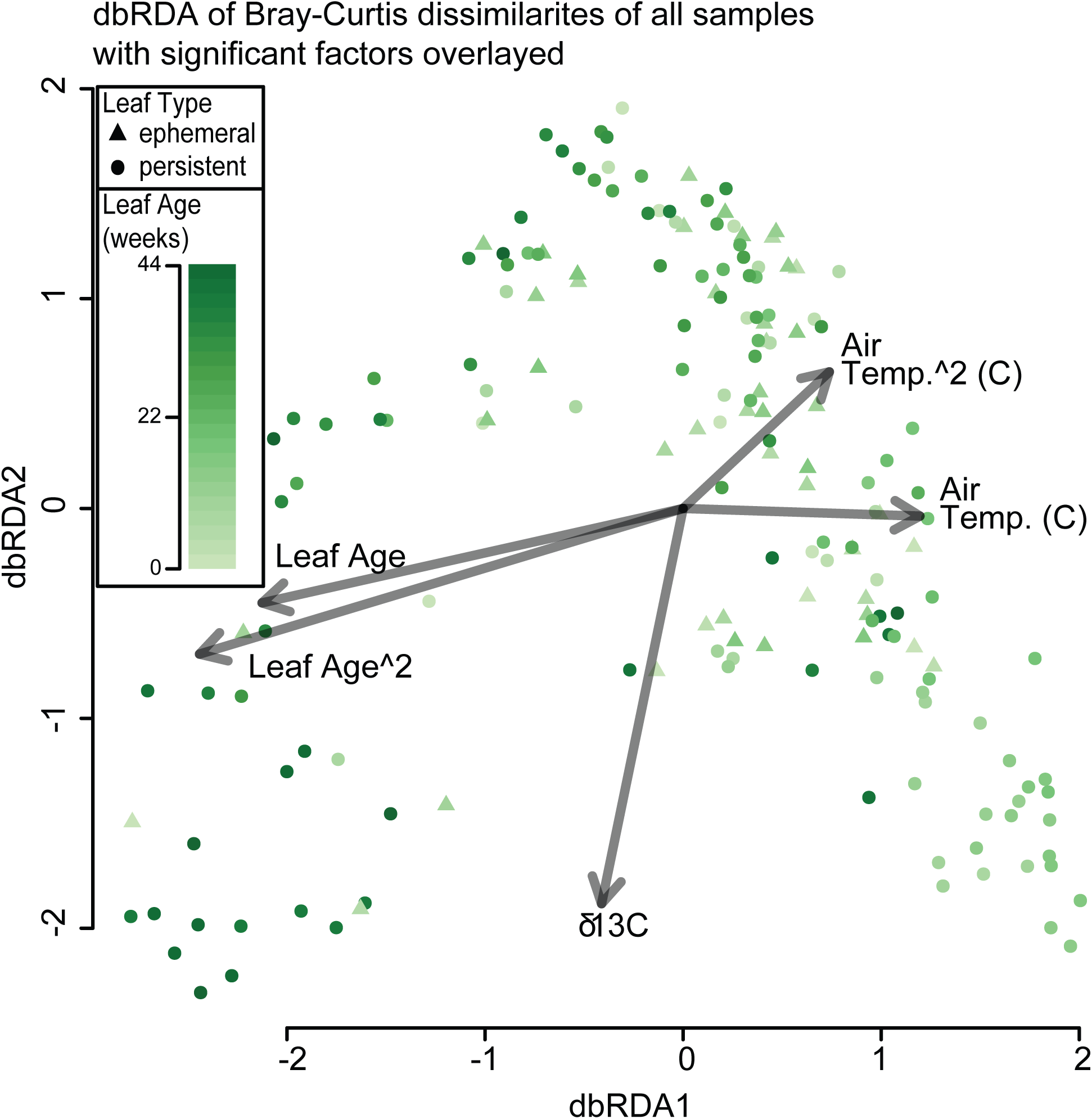
Beta diversity is correlated with leaf age, air temperature, δ^13^C, and leaf type. Distance-based Redundancy Analysis (dbRDA) ordination of Bray-Curtis dissimilarities among fungal communities (proportion explained = 0.123). Beta diversity (pairwise Bray-Curtis distance between communities) is significantly correlated with leaf age, leaf type, air temperature, and δ^13^C. Leaf age for individual samples is shown with a gradient of light to dark green, and leaf type is shown by triangles (ephemeral leaves) or circles (persistent leaves).

### Drivers of individual ASV presence and abundance

We found 20 genera that were present in at all sample dates: *Alternaria*, *Aureobasidium*, *Chaeothyrium*, *Chalara*, *Cladosporium*, *Comoclathris*, *Didymocyrtis*, *Endoconidioma*, *Filobasidium*, *Kabatina*, *Meristemomyces*, *Neocelosporium*, *Parafenestella*, *Penidiella*, *Phaeococcomyces*, *Preussia*, *Robertozyma*, *Staruosphaeria*, *Taphrina*, and *Thyrostroma* (Figure 5a). Of these only *Preussia* was found as an endophyte in our culture data, while the others were all epiphytes, or were not barcoded as part of our culture collection. Our sample on 06/03/2021 had an unusual 50% of relative abundance distributed among species that were not continually present (Figure 5a). The two weeks preceding this timepoint also experienced the largest relative increase in average air temperature (about 14°C, Figure 2b). This abnormal relative abundance was driven by two plants which had 92.5% and 57.6% relative abundance of non-continually present species. This change in relative abundance did not persist and the number of non-continually present species represented 1.5% of the total sequences at the next sampling date, 06/18/2021.

**Figure 5.**
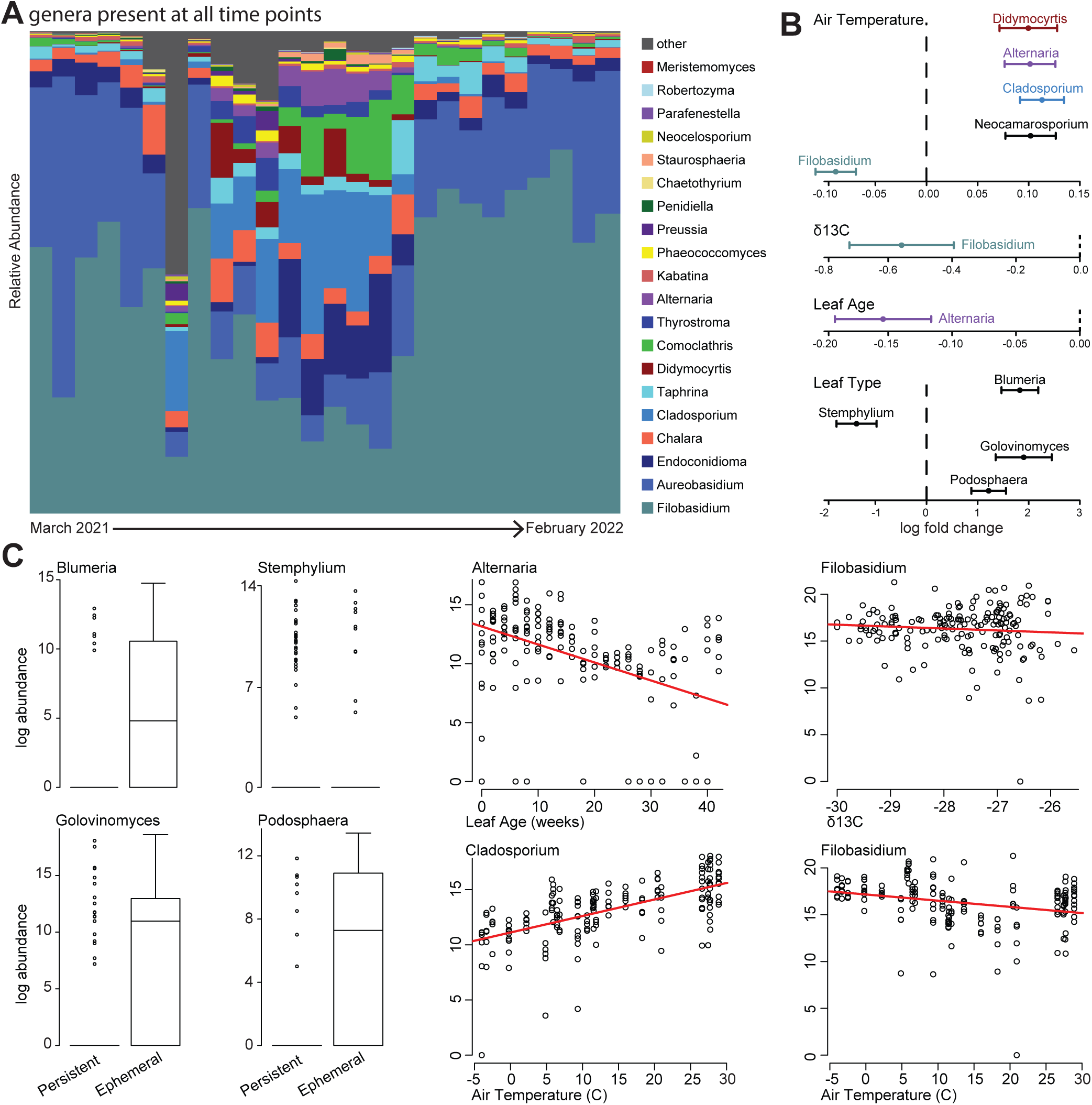
Factors influencing the abundances of different genera. (a) Relative abundances of the 20 persistent genera over time. All genera that were present in all time points were given a unique color; all others were binned as “other”. We observed that a few genera (e.g., *Filobasidium*, *Aureobasidium*, *Endoconidioma*, and *Chalara*) represented the majority of sequences at most timepoints. There was a more even distribution of taxa in the summer (center of the panel). (b) Results of ANCOMBC: log fold change from the baseline for each genus that we found to be significantly influenced by at least one factor. (c) Abundance of specific genera by variables with significant ANCOMBC results.

To further examine which factors influenced the different genera in the sagebrush phyllosphere, we estimated differential abundances using ANCOMBC. We found nine different genera (one basidiomycete and eight ascomycetes) that showed significant differential abundances across one or more factors. The majority (five) of these differentially abundant genera varied with temperature: *Filobasidium* (ANCOM-BC, logfold change (lfc) = -0.089, q = 5.059E-4), *Neocamarosporium* (lfc = 0.102, q = 0.003), *Cladosporium* (lfc = 0.113, q = 1.11E-05), *Alternaria* (lfc = 0.101, q = 0.003), and *Didymocyrtis* (lfc = 0.100, q = 0.029). Only one genus, *Alternaria*, varied with leaf age, and we found that it decreased with increasing leaf age (lfc = -.154, q = 0.003, Figures 5b and 5c). Four genera were differentially abundant on ephemeral leaves when compared to persistent (Figures 5b,c): *Podosphaera* (lfc = 1.216, q = 0.024), *Golovinomyces* (lfc = 1.902, q = 0.038), *Stemphylium* (lfc = -1.369, q = 0.035), and *Blumeria* (lfc = 1.826, q = 2.88E-05). We found one genus differentially abundant with δ13C, *Filobasidium* (lfc = -0.558, q = 0.048).

## Discussion

Here, we present a comprehensive and fine-scale characterization of a phyllosphere community over a full year. We used repeated measures to isolate the relative importance of weather, plant physiology, and substrate age, which tend to be collinear. To do this, we leveraged relevant features of sagebrush plants, which are evergreen and have a continuous succession of leaves as a substrate for microbial persistence. Most phyllosphere time-series studies up to this point have sampled over the growth season of plants with deciduous foliage and at irregular or less frequent time intervals (e.g., (Barge et al. 2019; Almario et al. 2022; Dove et al. 2021; VanWallendael et al. 2022; Wagner, Busby, and Balint Kurti 2020)). To our knowledge, this study represents the highest resolution longitudinal study of a phyllosphere community.

For all of our alpha diversity metrics, we observed large changes happening within two-week sampling windows (Figure 2c). These changes (e.g., a halving of the ENS from the end of October to the beginning of November) indicate that studies of phyllosphere communities using only single snapshots or sparse sampling will always be incomplete with regard to change over time, limiting conclusions about patterns and drivers of microbial species richness. Because changes in weather, host physiology, and leaf age are collinear in time, they may be confounded with other time-varying processes despite having independent effects on microbiome structure (Teller et al. 2016). Thus, we used a modelling approach that aimed to minimize bias in our estimation of independent effects of community age, climatic factors, and the host plant physiology on phyllosphere community diversity.

### Communities shift with substrate age, separate from weather and host physiology

Leaf age had the strongest effect on fungal richness. The hump-shaped effect we saw for leaf age (separate from leaf type; Figure 3a), is likely due to successional processes: species accumulate on a new leaf (a novel community substrate) with gradual dispersal, and then species numbers decline as either competition or leaf senescence create a harsher environment. Across studies, there is no clear consensus of how alpha diversity changes with leaf age. In the *Arabidopsis thaliana* phyllosphere, Beilsmith et al. (Beilsmith, Perisin, and Bergelson 2021) reported an increase in bacterial Shannon diversity throughout the growing season while Almario et al. (Almario et al. 2022) reported no change for the bacterial community and a decrease in the fungal and oomycete communities. Mixed results have also been reported for various crop species (Wagner, Busby, and Balint Kurti 2020; Copeland et al. 2015; Grady et al. 2019). Surprisingly, in our study total fungal abundance showed a very different pattern, with an initial slight decrease and then a strong increase on leaves more than 30 weeks old (Figure 3A). Together with the richness data, this suggests that certain taxa multiply on older leaves, perhaps because they are strong competitors in the habitat, or because they are beginning to degrade the leaves. We noticed that a few taxa (e.g. *Filobasidium sp.*, *Aureobasidium sp.*) proliferated when richness was low. Foliar microbes have been found to contribute to decomposition of leaf litter (Wolfe and Ballhorn 2020), including several highly abundant taxa found in our study (e.g. *Aureobasidium, Cladosporium*) and the proliferation of these taxa may be linked to the senescence and decomposition of leaves at later life stages.

Separate from leaf age, we also found that sagebrush’s ephemeral leaves tended to have higher abundance than persistent leaves. Ephemeral leaves have lower levels of secondary chemicals (e.g. monoterpenes, phenols) and higher levels of crude proteins and water as well as more surface area compared to persistent leaves (Campos et al. 2011). These conditions may provide a more favorable substrate for fungal proliferation despite the fact that ephemeral leaves exist during the hottest time of the year.

### Communities shift with weather conditions, separate from substrate age and host physiology

We also observed changes in phyllosphere communities due to weather. Air temperature can select for fungal species due to species-level differences in survivability and proliferation at different temperatures. For example, our two most abundant genera, *Filobasidium* and *Aureobasidium*, were more common in colder times of year (Figure 5a) and both contain species that exhibit physiological activity in extreme cold (Zalar and Gunde-Cimerman 2014), producing glycerol during osmotic stress (Chi et al. 2022; Tekolo et al. 2010). A higher ENS, and more even distribution of taxa, was observed in the summer months (Figures 2c, 5a). We observed that ENS increases with temperature to a point (Figure 3b) indicating that the most taxa were present in this system at about 28°C. Abundance was strongly affected by precipitation. Precipitation is considered a dispersal agent of new microbes into the phyllosphere (Mechan Llontop et al. 2021), but it did not have a clear effect on species richness in our study. However, the positive relationship between abundance and precipitation observed in this analysis corroborates how precipitation can selectively increase the abundance of some fungal taxa in both the leaf and soil microbiome (Chen et al. 2021; Herrera et al. 2011). In our system, the increase in fungal abundance with precipitation was dominated by *Filobasidium spp.* and is likely due to the fact that water is limiting in the cold desert conditions in the sagebrush steppe ecosystems. Water is essential for fungal metabolism and reproduction and tolerance of osmotic conditions is different between fungal species (Duran, Cary, and Calvo 2010). In yeasts, the high osmolarity glycerol pathway is often activated during osmotic stress, providing for the production of glycerol and increased osmotic pressure in the cell (Davis, Burlak, and Nicholas 2000).

### Evergreen leaves are a reservoir of persistent microbial species

In this study, we present the first characterization of the sagebrush phyllosphere community, unique from the plant perspective because of sagebrush’s chemical profile and status as foundation species in a geographically extensive ecosystem. Most phyllosphere community studies of evergreen plants have been focused on microbial functions in coniferous plants, including nitrogen fixation (Shi et al. 2023), litter decomposition (Sun et al. 2021), and the interactions of microbes with defoliating insects (Beule et al. 2017). No studies, to our knowledge, have examined the continual persistence of microbes in an evergreen phyllosphere, a unique opportunity for specialization of persistent microbes to a single host species and even potential co-evolution between the host and its persistent microbes. We observed the continual persistence of 20 genera of fungi (Figure 5a), which, theoretically, could remain present on sagebrush leaf substrates indefinitely. Further research is needed to determine if this pattern reflected persistence of specific strains, and whether persistent strains are locally adapted to their leaf environments. Evergreen plants are often foundation species in their ecosystems and also are reservoirs of potential leaf specialist species which may disperse onto annual foliage of other plant species. The existence of such reservoirs adds another dimension to interspecies interactions in ecosystems and may augment the importance of already essential plant species.

### Taxa-level shifts in relative abundance

Decrypting the function of leaf communities is an ongoing project for phyllosphere ecologists. A first step is modeling the response of individual taxa to temporally collinear variables. Here we have the first understanding of this in the sagebrush system. Overall, we observed 25% Basidiomycetes and 75% Ascomycetes, which corroborates the dominance of the phylum Ascomycota in other studies of phyllosphere communities (Chen et al. 2020). Over the course of a year in this one plant subspecies, we observed 18 classes, 62 orders, 165 families, and 365 genera. Understanding the responses of taxa to environmental variables can help predict community composition [56], and can lead to a better understanding of different species’ ecological roles. The genera we detected cover a range of potential functions, from pathogenic to beneficial (Răut et al. 2021).

## Conclusions

The leaf as a microbial community substrate is bounded in time by budding and senescence. Plant species at any moment in time are highly variable in the presence or absence and age of their leaves. Thus, time-series analyses of leaf microbiomes are essential and yet complex. Our study, unique for the combination of its resolution and length, provides novel data of the change in leaf communities over time. We recorded rapid shifts in community diversity in short periods of time in response to changes in weather and host plant physiology, highlighting the necessity of repeated regular sampling at short time intervals. Individual taxa responded differently to our measured factors and twenty of the over three hundred genera recorded were present at all the time points, demonstrating that sagebrush plants harbor a core persistent phyllosphere microbiome with the potential for long-term coevolution. Evergreen plants can represent continuous reservoirs of highly adapted microbial species and may serve as sources of specialist foliar microbes for deciduous plant species. For a comprehensive understanding of the phyllosphere microbiome and its role in ecosystem functioning, further research is needed to characterize specialized species living on evergreen plants that may have evolved unique functions to fit the phyllosphere niche space.

## Acknowledgements

This research was supported by a seed grant from the NSF Idaho EPSCoR Program and by the National Science Foundation under the GEM3 award number OIA-1757324. Next-Generation Sequencing was performed by the Molecular Research Core Facility at Idaho State University (RRID:SCR_012598). This equipment was made possible by NIH/NCRR Award #P20RR016454. Special thanks to the members of the Bittleston Lab for assistance with data gathering and lab work and to Jennifer Forbey, Marcelo Serpe, and Posy Busby.

