## Supplement 1 for "Weather and leaf age separately contribute to temporal shifts in phyllosphere community structure and composition"

Appendix S1

Journal: Ecosphere

### Title: Weather and leaf age separately contribute to temporal shifts in phyllosphere community structure and composition

Authors: Jacob A. Heil[1]*, Allison Simler-Williamson[1], Miranda L. Striluk[2], Danielle Trawick[3], Rachel Capezza[1], Chadwick DeFehr[1], Aubrey Osorio[1], Bruce Finney[3], Kathryn G. Turner[3], Leonora S. Bittleston[1]

#### Supplemental Methods

##### *Leaf Types*

Sagebrush plants maintain functioning foliage throughout the year and have different leaf types depending on the time of year. When we collected our samples, we made sure to differentiate between two different leaf types: persistent and ephemeral (Figure 2a). For identification in the field, we characterized persistent leaves as being relatively smaller than ephemeral leaves, having three relatively small and regular lobes at the tip of the leaf, and being present on the plant from the middle of summer to the middle of the next spring (McDonough, Harniss, and Campbell 1975; Miller and Shultz 1987). To quantify this difference, we calculated the area of ten representative leaves from each type and found an average size of .345 $cm^2$ for persistent leaves and .671 $cm^2$ for ephemeral. We collected persistent leaves from March 12, 2021 (our first sample date) to April 9, 2021 after which we did not observe any persistent leaves of reasonable size on our plants until we began collecting them again on July 2, 2021 and continued until the end of the study on February 24, 2022. We collected persistent leaves on a total of 22 sample dates. We characterized ephemeral leaves as being relatively larger than persistent leaves, having three or more relatively large and irregular lobes at the tip of the leaf, and being present on the plant beginning in the middle of spring until the late summer. We collected ephemeral leaves from May 22, 2021 when we first observed their presence, to July 30, 2021 when we no longer observed any on the plants. During four sampling dates we collected both leaf types and processed them separately.

*Microbial Culturing*

For our culturing data set we collected six leaves at each time point in sterile tubes, placed them on ice in the field, and transported them to the lab where we cultured them on the same day as sampling. We cultured both the leaf epiphytes and endophytes on 60 x 15mm potato dextrose agar (PDA) petri dishes. To culture leaf epiphytes, we added 200 µL of autoclaved, deionized (DI) water to each of our six leaf tubes in a biosafety cabinet and vortexed the tube for two minutes before placing the leaf wash water on a petri dish with autoclaved glass beads and manually agitating the petri dishes to spread the leaf wash evenly across the surface of the media. To culture leaf endophytes, we took the same leaves that we had just washed for epiphytes and cut them into 1mm x 1mm squares, placed them in metal mesh tea infusers and surface sterilized the leaves by washing them in 8.5% tween, 70% Ethanol, and two separate autoclaved DI water baths for two minutes each. After the leaves had been surface sterilized, they were transferred to a PDA plate. All epiphytic and endophytic cultures were allowed to grow for one week at room temperature. After one week, we collected community data from the leaf communities by counting all unique morphospecies and their abundances from each plant at each time point. We pooled all communities at the plant level including merging the epiphyte and endophyte communities into one data set. In total we collected 180 different sample communities. We subcultured all visibly unique morphospecies from each plant for further DNA barcoding and developed a library of isolated morphospecies. We extracted DNA from pure isolates using the Qiagen DNeasy Powerlyzer Microbial kit (Cat # 12255-50). We amplified DNA barcodes from our isolates using the 27F and 1492R primers for suspected bacteria and ITS1f and ITS4 primers for suspected fungi. The amplified DNA was cleaned and sent to GENEWIZ (Azenta) for Sanger sequencing.

*Taxonomy Assignment*

We used the R package *rBLAST* (Hahsler and Nagar 2024) to assign taxonomy to the ASVs. We built a database of our DNA sequences generated from our barcoding of selected sagebrush leaf fungal cultures. This database was not as comprehensive as ITS RefSeq, and with it we matched about 59% of our sequences to the species level. To assign taxonomy, we first matched our sequences to this local sagebrush database and accepted any exact matches and then matched our ASV sequences to the most similar matches in NCBI's ITS RefSeq Fungi database (O'Leary et al., 2016). For each sequence in the FASTA file we selected the matches with the highest Bit score values, translated their GenBank Accession IDs into NCBI Taxonomy IDs using the *taxonomizr* package (Sherrill-Mix 2023), and built a table of their full taxonomy using the *taxize* package (Chamberlain et al. 2022). We then compared these matches for conflicting results at each taxonomic level. For each level we accepted the most likely assignment (highest Bit score). If there were multiple conflicting most likely assignments, we assigned it "NA". We also ran this consensus process with other databases other than NCBI's ITS RefSeq. We found that there was higher species resolution with RefSeq (88%) compared to UNITE (85%; (Abarenkov et al. 2023)).

*Change in alpha diversity with time*

To determine if there was significant variation between sample dates we fit negative binomial glmms with each of our response variables (ASV Richness, Effective Number of Species, Total Abundance), sampling date as the only predictor, and Plant ID as a random intercept (Figure 2). We observed significant differences between sampling dates, indicating a significant change in time. We developed another model structure to determine how factors that are collinear in time are influencing change in our samples. We developed Bayesian glmms to determine the effect of variables on alpha diversity. For or response variables (ASV richness, total abundance, and effective number of species) using the brms package and weakly informative priors (Bürkner 2017). Each model contained: 1) leaf age (in days, measured as time since first collection) to describe shifts in community composition associated with substrate age and colonization; 2) variables describing mean air temperature, mean wind speed, and total precipitation for the two week interval preceding each sampling point, to describe the effects of various weather drivers; and 3) leaf $\delta^{13}C$ and $\delta^{15}N$, to describe seasonal variation in host physiology, associated with photosynthetic activity and drought stress. We included a quadratic term for mean air temperature and leaf age, to model possible upper thermal limits for fungi and hump-shaped relationships between time and diversity, respectively. Models also included a varying intercept for plant identity, to account non-independence of repeated measures in its error structure. All continuous variables were centered and scaled by two standard deviations.

Supplemental Figures

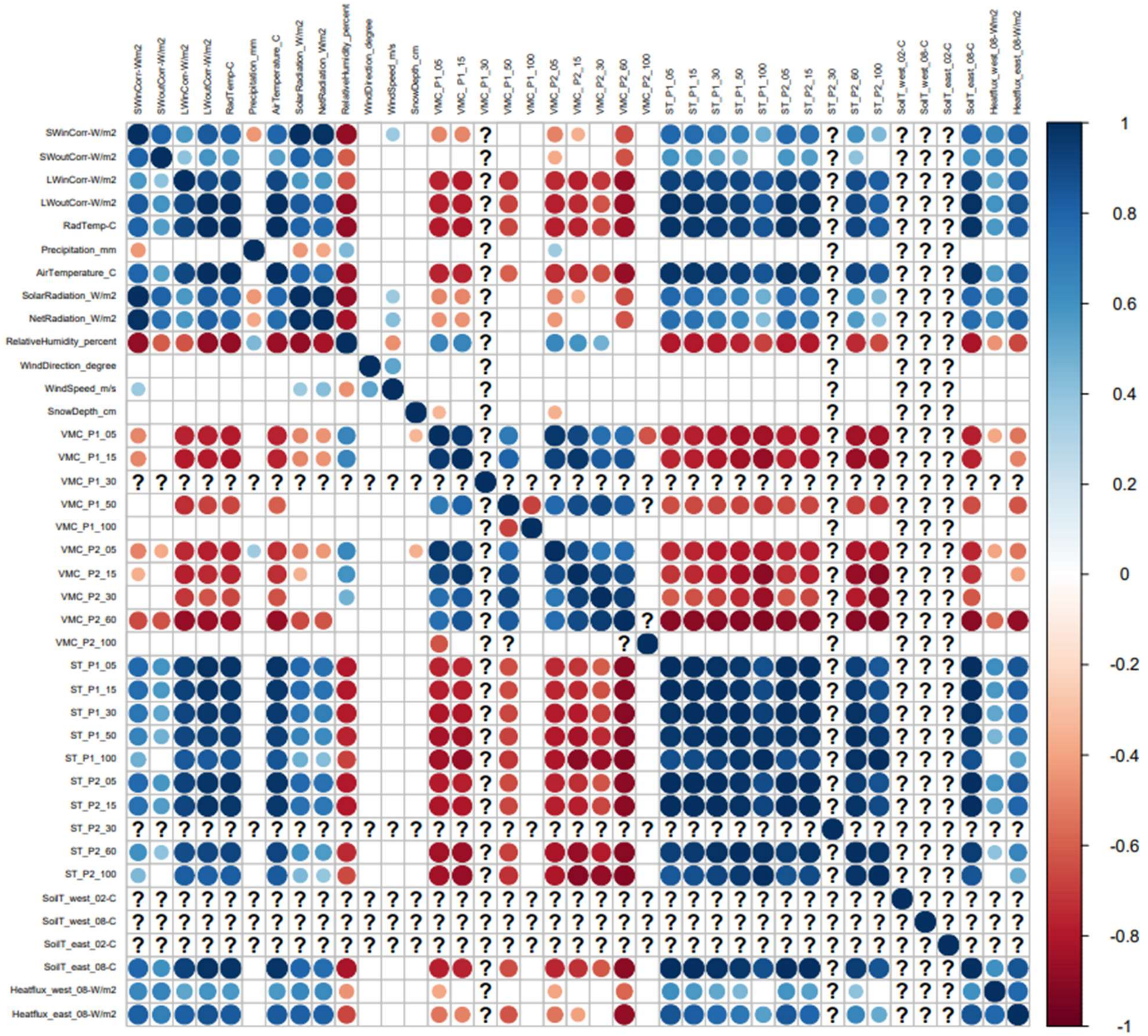


**Figure S1. Correlation plot of environmental variables**. Pairwise Pearson correlations for all environmental variables averaged over 10-day (240 hour) periods. Darker blue indicates higher correlation, darker red indicates higher anticorrelation, and question marks indicate incomplete data. SW = shortwave radiation. LW = longwave radiation. Radtemp = radiometer temperature. VMC = volumetric moisture content. We found that Air Temperature is strongly correlated or anti-correlated with the majority of other factors and for this reason used it in our model. We used precipitation in our model because it was not correlated with air temperature and contributes to the survivability and dispersal of microorganisms. We used wind speed in our model because it is not correlated with air temperature and contributes to the dispersal of microorganisms. ST and SoilT = soil temperature.

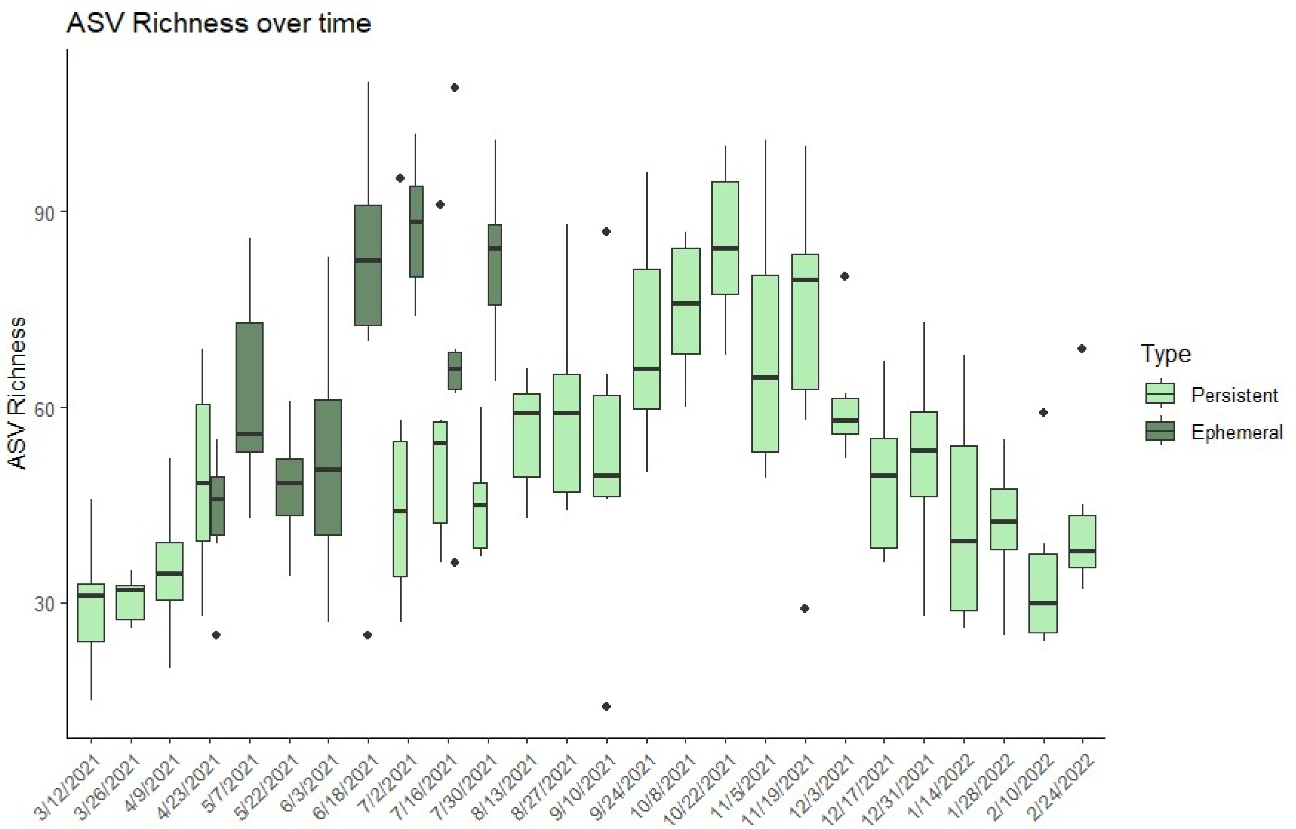
ASV Richness over time
ASV Richness
90
60
30
3/12/2021
3/26/2021
4/9/2021
4/23/2021
5/7/2021
5/22/2021
6/3/2021
6/18/2021
7/2/2021
7/16/2021
7/30/2021
8/13/2021
8/27/2021
9/10/2021
9/24/2021
10/8/2021
10/22/2021
11/5/2021
11/19/2021
12/3/2021
12/17/2021
12/31/2021
1/14/2022
1/28/2022
2/10/2022
2/24/2022
Type
Persistent
Ephemeral

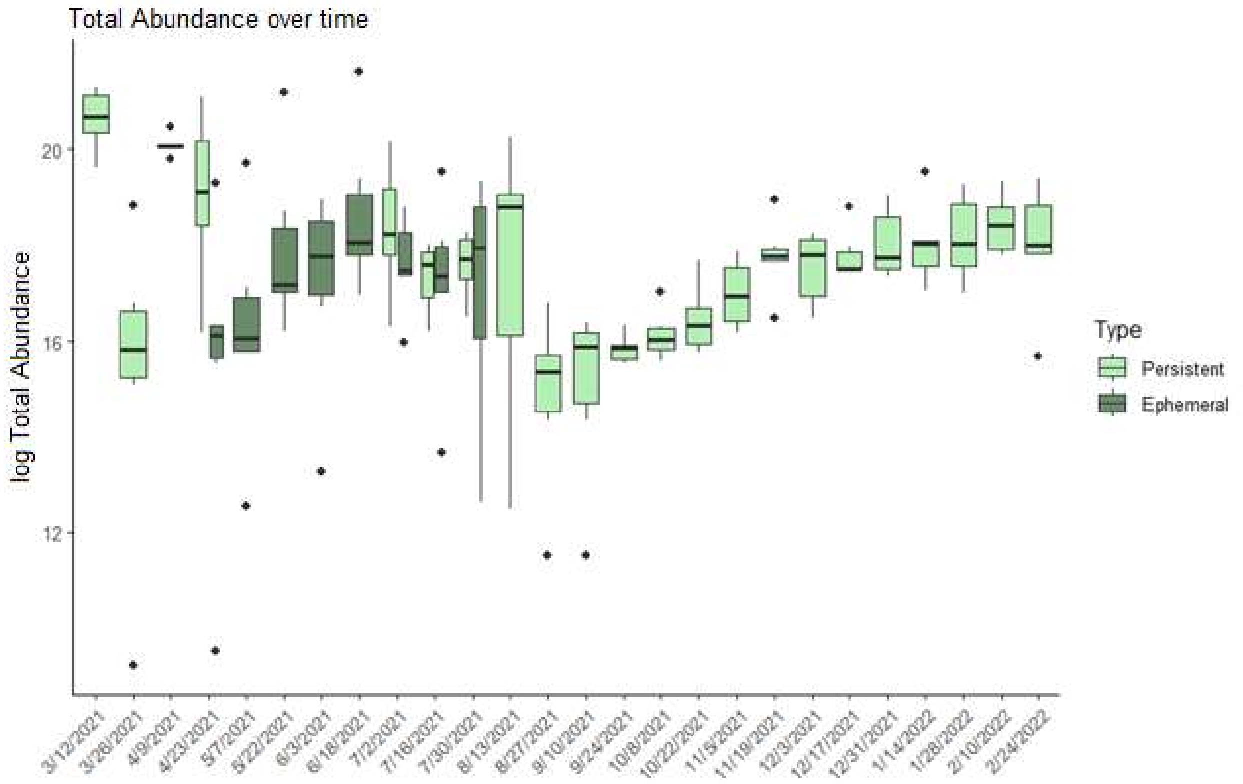
Total Abundance over time
log Total Abundance
20
16
12
3/12/2021
3/26/2021
4/9/2021
4/23/2021
5/7/2021
5/22/2021
6/3/2021
6/18/2021
7/2/2021
7/16/2021
7/30/2021
8/13/2021
8/27/2021
9/10/2021
9/24/2021
10/8/2021
10/22/2021
11/5/2021
11/19/2021
12/3/2021
12/17/2021
12/31/2021
1/14/2022
1/28/2022
2/10/2022
2/24/2022
Type
Persistent
Ephemeral

**Figure S2. ASV richness and total abundance over time.** We found significant changes in both ASV richness and total fungal abundance over time. Boxes are colored by leaf type.

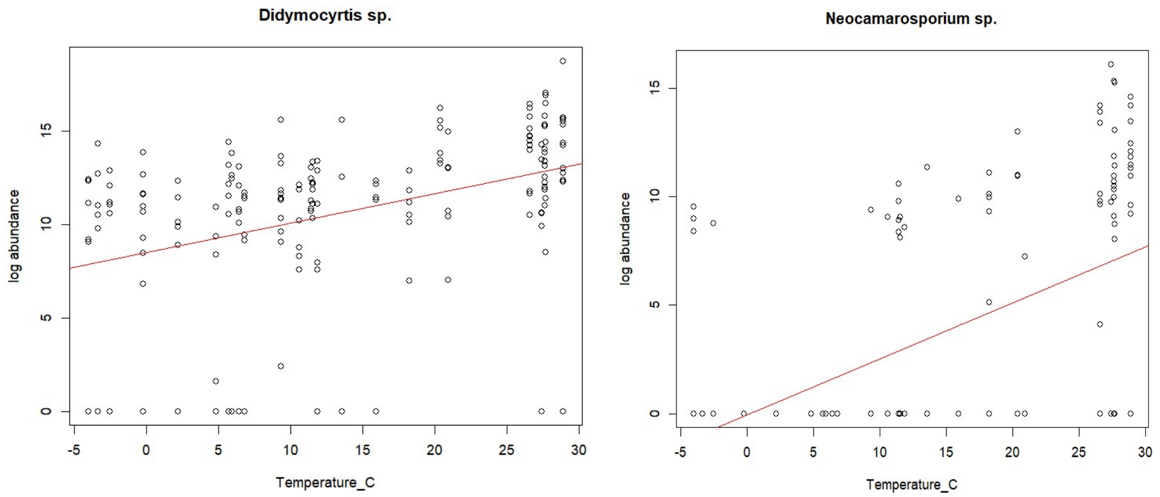


**Figure S3. More ANCOMBC results.** *Didymocyrtis sp.* abundance was positively correlated with air temperature (lfc = -1.181, p = 0.0162) and *Neocamarosporium sp.* abundance was as well (lfc = 0.234, p < 0.001).

Supplemental Tables

| | | Deviance | Resid. Df | Resid. Dev. | Pr(>Chi) |
|---|---|---|---|---|---|
| ASV richness | **Date** | 1.39E+02 | 153 | 181.7182 | 7.14E-18* |
| | **1 \| Plant** | 2.27E-13 | 153 | 181.7182 | *NA* |
| ASV abundance | **Date** | 1.95E+02 | 153 | 212.3285 | 2.82E-28* |
| | **1 \| Plant** | 1.82E-06 | 153 | 212.3285 | *NA* |
| effective number of species | **Date** | 2.73E+02 | 153 | 162.6398 | 1.35E-43* |
| | **1 \| Plant** | 2.84E-14 | 153 | 162.6398 | *NA* |

**Table S1. ANOVA of model results.** For each of our response variables we tested the effect of sample date using a generalized linear model with a negative binomial distribution and individual plants as random intercepts. To test significance of difference between results in the model we used an ANOVA, the results of which are presented in this table. We found that there were significant differences between sample dates in all models.

*Results for ASV richness model: richness.ITS ~ scale(LeafAge) + scale(I(LeafAge^2)) + Type + scale(AirTemperature_C) + scale(I(AirTemperature_C^2)) + scale(Precipitation_mm) + scale(WindSpeed_m.s) + scale(Delta15N) + scale(Delta13C) + (1 | Plant)*

| | Estimate | Est.Error | l-95% CI | u-95% CI | Rhat | Bulk_ESS | Tail_ESS |
|---|---|---|---|---|---|---|---|
| **Intercept** | 4.05 | 0.09 | 3.87 | 4.24 | 1 | 8247 | 9260 |
| **scaleLeafAge** | 0.66 | 0.11 | 0.45 | 0.87 | 1 | 13035 | 13575 |
| **scaleILeafAgeE2** | -0.69 | 0.09 | -0.87 | -0.5 | 1 | 12955 | 13331 |
| **TypePersistent** | -0.08 | 0.07 | -0.21 | 0.05 | 1 | 22001 | 15959 |
| **scaleAirTemperature_C** | 0.34 | 0.1 | 0.15 | 0.53 | 1 | 11204 | 13010 |
| **scaleIAirTemperature_CE2** | -0.22 | 0.1 | -0.41 | -0.03 | 1 | 10364 | 13008 |
| **scalePrecipitation_mm** | -0.01 | 0.03 | -0.06 | 0.04 | 1 | 24428 | 15071 |
| **scaleWindSpeed_m.s** | 0.03 | 0.03 | -0.03 | 0.1 | 1 | 15448 | 15534 |
| **scaleδ15N** | 0.01 | 0.03 | -0.05 | 0.06 | 1 | 21006 | 13951 |
| **scaleδ13C** | -0.02 | 0.04 | -0.09 | 0.06 | 1 | 15592 | 15716 |

*Results for total abundance model: abundance.ITS ~ scale(LeafAge) + scale(I(LeafAge^2)) + Type + scale(AirTemperature_C) + scale(I(AirTemperature_C^2)) + scale(Precipitation_mm) + scale(WindSpeed_m.s) + scale(Delta15N) + scale(Delta13C) + (1 | Plant)*

| | Estimate | Est.Error | l-95% CI | u-95% CI | Rhat | Bulk_ESS | Tail_ESS |
|---|---|---|---|---|---|---|---|
| **Intercept** | 18.68 | 0.35 | 17.98 | 19.36 | 1 | 18119 | 20697 |
| **scaleLeafAge** | -1.16 | 0.46 | -2.05 | -0.25 | 1 | 22049 | 25762 |
| **scaleILeafAgeE2** | 1.96 | 0.4 | 1.17 | 2.75 | 1 | 21737 | 25763 |
| **TypePersistent** | -0.65 | 0.28 | -1.2 | -0.1 | 1 | 33318 | 31219 |
| **scaleAirTemperature_C** | -0.29 | 0.42 | -1.13 | 0.52 | 1 | 20879 | 26601 |
| **scaleIAirTemperature_CE2** | 0.54 | 0.41 | -0.26 | 1.35 | 1 | 19297 | 24303 |
| **scalePrecipitation_mm** | 0.29 | 0.11 | 0.09 | 0.51 | 1 | 36227 | 29370 |
| **scaleWindSpeed_m.s** | -0.17 | 0.14 | -0.44 | 0.1 | 1 | 32636 | 29700 |
| **scaleδ15N** | -0.07 | 0.12 | -0.31 | 0.17 | 1 | 28482 | 29163 |
| **scaleδ13C** | 0.04 | 0.17 | -0.29 | 0.37 | 1 | 26264 | 29363 |

*Results for effective number of species model: effective.ITS ~ scale(LeafAge) + scale(I(LeafAge^2)) + Type + scale(AirTemperature_C) + scale(I(AirTemperature_C^2)) + scale(Precipitation_mm) + scale(WindSpeed_m.s) + scale(Delta15N) + scale(Delta13C) + (1 | Plant)*

| | Estimate | Est.Error | l-95% CI | u-95% CI | Rhat | Bulk_ESS | Tail_ESS |
|---|---|---|---|---|---|---|---|
| **Intercept** | 1.97 | 0.14 | 1.69 | 2.26 | 1 | 15694 | 16370 |
| **scaleLeafAge** | 0.4 | 0.16 | 0.08 | 0.72 | 1 | 25331 | 26372 |
| **scaleILeafAgeE2** | -0.59 | 0.15 | -0.89 | -0.29 | 1 | 24686 | 27255 |
| **TypePersistent** | 0.08 | 0.1 | -0.11 | 0.27 | 1 | 39525 | 31200 |
| **scaleAirTemperature_C** | 0.78 | 0.17 | 0.45 | 1.11 | 1 | 25738 | 28040 |
| **scaleIAirTemperature_CE2** | -0.38 | 0.16 | -0.69 | -0.06 | 1 | 23611 | 26454 |
| **scalePrecipitation_mm** | 0.07 | 0.04 | -0.01 | 0.15 | 1 | 45675 | 29881 |

| | | | | | | | |
|---|---|---|---|---|---|---|---|
| **scaleWindSpeed_m.s** | -0.02 | 0.05 | -0.12 | 0.09 | 1 | 30984 | 29682 |
| **scaleδ15N** | 0.01 | 0.04 | -0.07 | 0.1 | 1 | 39041 | 31036 |
| **scaleδ13C** | 0.07 | 0.06 | -0.04 | 0.19 | 1 | 30146 | 30192 |

**Table S2. Model Results.** We modeled our three response variables with the explanatory variables leaf age, leaf type, air temperature, precipitation, wind speed, δ15N, δ13C, and individual plants as random intercepts.

```
Permutation test for dbrda under reduced model
Marginal effects of terms
Permutation: free
Number of permutations: 999

Model: dbrda(formula = dist.ITS ~ scale(LeafAge) + scale(I(LeafAge^2)) + Type
+ scale(AirTemperature_C) + scale(I(AirTemperature_C^2)) + scale(Precipitatio
n_mm) + scale(WindSpeed_m.s) + scale(Delta15N) + scale(Delta13C), data = meta
data_plant, distance = "bray", na.action = na.exclude, Condition = metadata_p
lant$Plant)
                              Df SumOfSqs      F Pr(>F)    
scale(LeafAge)                 1    0.629 1.6382  0.007 ** 
scale(I(LeafAge^2))            1    1.207 3.1452  0.001 ***
Type                           1    0.618 1.6088  0.013 *  
scale(AirTemperature_C)        1    1.053 2.7431  0.001 ***
scale(I(AirTemperature_C^2))   1    0.923 2.4044  0.003 ** 
scale(Precipitation_mm)        1    0.377 0.9822  0.452    
scale(WindSpeed_m.s)           1    0.399 1.0400  0.334    
scale(Delta15N)                1    0.440 1.1461  0.192    
scale(Delta13C)                1    0.830 2.1622  0.001 ***
Residual                     167   64.107                  
```

**Table S3. Permutation test for significance for our dbRDA.** We calculated Bray-Curtis dissimilarity of our data and modeled community dissimilarity a response variable using dbRDA. This beta diversity model adheres as closely as possible in structure to our alpha diversity model. This table contains the results of an ANOVA-like permutation test run on the dbRDA model result.

Bürkner, Paul-Christian. 2017. “Brms: An R Package for Bayesian Multilevel Models Using Stan.”

Chamberlain, Scott, Eduard Szoecs, Zachary Foster, Zebulun Arendsee, Carl Boettiger, Karthik Ram, Ignasi Bartomeus, et al. 2022. “Taxize: Taxonomic Information from Around the Web.” https://cran.r-project.org/web/packages/taxize/index.html.

Hahsler, Michael, and Anurag Nagar. 2024. “rBLAST: R Interface for the Basic Local Alignment Search Tool.” http://bioconductor.org/packages/rBLAST/.

McDonough, W.T., R.O. Harniss, and R.B. Campbell. 1975. “Morphology of Ephemeral and Persistent Leaves of Three Subspecies of Big Sagebrush Grown in a Uniform Environment.” *Great Basin Naturalist* 35 (3): 10.

Miller, Richard F., and Leila M. Shultz. 1987. “Development and Longevity of Ephemeral and Perennial Leaves on Artemisia Tridentata Nutt. Ssp. Wyomingensis.” *The Great Basin Naturalist*, 227–30.

Sherrill-Mix, S. 2023. “Taxonomizr: Functions to Work with NCBI Accessions and Taxonomy_.” https://CRAN.R-project.org/package=taxonomizr.
